# Mena-dependent local translation of PI3K-p85 coordinates the axonal regenerative program upon injury

**DOI:** 10.1101/2025.09.29.679150

**Authors:** Sofia Pasadaki, Nikoletta Triantopoulou, Christina Koupepia, Eirini Triantafyllidou, Eirini Kokolaki, Martina Samiotaki, Marina Vidaki

## Abstract

Local translation (LT) is one of the first processes activated after nerve injury and is critically coupled with the intrinsic ability of axons to grow and regenerate. However, regulation of LT in adult axons remains highly unexplored. Here we identify the actin regulator Mena, as a key mediator of adult axon regeneration, via its control of local protein synthesis. We show that Mena directly associates with PI3K-p85 mRNA and regulates its LT in injured sciatic nerve axons, thereby activating the downstream AKT/mTOR pathway. Genetic ablation of Mena disrupts this intrinsic response, resulting in impaired axonal LT and diminished axon regeneration *in vivo*. Our findings identify a Mena-dependent mechanism that remodels the local axonal proteome and reveal a critical link between mRNA regulation and metabolic signaling during axon regeneration.

## Introduction

During development neuronal axons receive continuous guidance signals from their environment, as they travel long distances to reach their synaptic targets, and mature axons depend on external cues for maintenance and repair. In either case, axons must quickly process environmental information, often independently of the soma. Local mRNA translation (LT) is a key mechanism for this autonomy, and disruption of the process during development is implicated in aberrant axon guidance, defective synaptic function, and neurodevelopmental disorders(Verma et al. 2005; Jung et al. 2012; Baleriola et al. 2014). Increasing evidence indicates that axonal LT continues to play roles in mature axons and injury-induced axon regeneration in particular(Verma et al. 2005; Jung et al. 2012; Baleriola et al. 2014). It is one of the first processes activated after injury and numerous studies demonstrate that enhancing LT can significantly improve the regenerative capacity of axons, both *in vitro* and *in vivo*(Saxton and Sabatini 2017; Christie et al. 2010), whereas blocking LT attenuates regeneration of severed axons in culture(Verma et al. 2005) and after nerve injury *in vivo*(Donnelly et al. 2011). Thus, the regenerative capacity of axons is highly correlated to their intrinsic ability to synthesize proteins *in situ*, and modulate their proteome on demand. Despite the evident role of LT in axon regeneration, relatively little is known regarding its regulation in adult axons, and most of the existing knowledge relies on the identification of specific mRNAs found in axons, rather than the mechanisms that regulate their LT, and how they might be coupled to injury signals.

Previous research has identified a ribonucleoprotein complex (RNP) that regulates LT in developing cortical axons(Vidaki et al. 2017). A key component of the complex is Mena, which is a member of the Ena/VASP family of proteins that promote actin polymerization and regulate cell adhesion and motility(Gertler et al. 1996; Krause et al. 2003; Drees and Gertler 2008; Gupton et al. 2012; Bear et al. 2002). Mena, along with VASP and EVL, the other two members of the family, are essential for axon guidance and growth cone dynamics(Dent et al. 2011; Kwiatkowski et al. 2007; McConnell et al. 2016), and Mena/VASP-null, or Mena/VASP-EVL- triple null neurons fail to respond to major guidance cues like Netrin(Lebrand et al. 2004) and Slit(McConnell et al. 2016; Bashaw et al. 2000). Moreover, triple-null mice exhibit severe axon guidance defects in their nervous system(McConnell et al. 2016; Lanier et al. 1999), while single Mena-deficiency leads to commissural axon midline crossing phenotypes with low penetrance(Lanier et al. 1999). Although those studies reveal the importance of Ena/VASP proteins in axon development, they also show that the actin-regulating function of Mena can be partially compensated in neurons by other members of the Ena/VASP family (Kwiatkowski et al. 2007; Lebrand et al. 2004; Menzies 2004; Lanier et al. 1999). However, in a separate mode of function, Mena was found to interact with RNA-binding proteins (RBPs) that are known translational regulators (e.g. HnrnpK and PCBP1(Gebauer and Hentze 2004)) and cytosolic mRNAs, forming a ribonucleoprotein complex (Mena-RNP)(Vidaki et al. 2017). This property was identified to be unique to Mena, and no other members of the Ena/VASP family, and it was demonstrated that Mena is necessary for LT of the Mena-RNP-associated mRNAs in developing axons(Vidaki et al. 2017). Although Mena has been extensively studied during nervous system development, its function in adult axons has not been established. However, its distinctive characteristics, render it an ideal candidate molecule to moderate adult axon behavior, particularly with respect to growth and regeneration after injury.

In this study, we identify Mena as a major regulator of axonal LT in adult peripheral nerves. We show that Mena couples injury signals to the synthesis of PI3K-p85 subunit, via association with the *Pik3r1* mRNA, leading to the activation of the PI3K/Akt/mTOR pathway. Loss of Mena in mice abolishes this LT switch leading to impaired mTOR auto-regulation, compromised activation of translation effectors like S6K1, S6 and eEF2, and ultimately a dramatic decrease of axonal protein synthesis. At the functional level, Mena-deficient axons fail to regenerate *in vivo*, after sciatic nerve injury. Taken together, our data uncover a previously unrecognized mechanism that links actin-associated proteins to translational control in response to injury, revealing the central role that Mena has in mediating the intrinsic ability of axons to regenerate.

## Results

### Mena- deficiency affects the regenerative capacity of adult SN axons after injury *in vivo*

To explore the function of Mena in adult axons, we first studied its expression levels and pattern in adult mouse SNs (2month old, 2MO) *in vivo*. Using western blot and quantitative PCR (qPCR) from whole nerve protein and mRNA extracts respectively, we found that Mena is expressed in axons of the adult SN (Fig. S1E, F), while immunofluorescence (IF) and confocal microscopy on longitudinal cryosections from isolated SNs revealed localization and distribution of the protein along SN axons (Fig. 1A, Fig. S1A).

**Figure 1:**
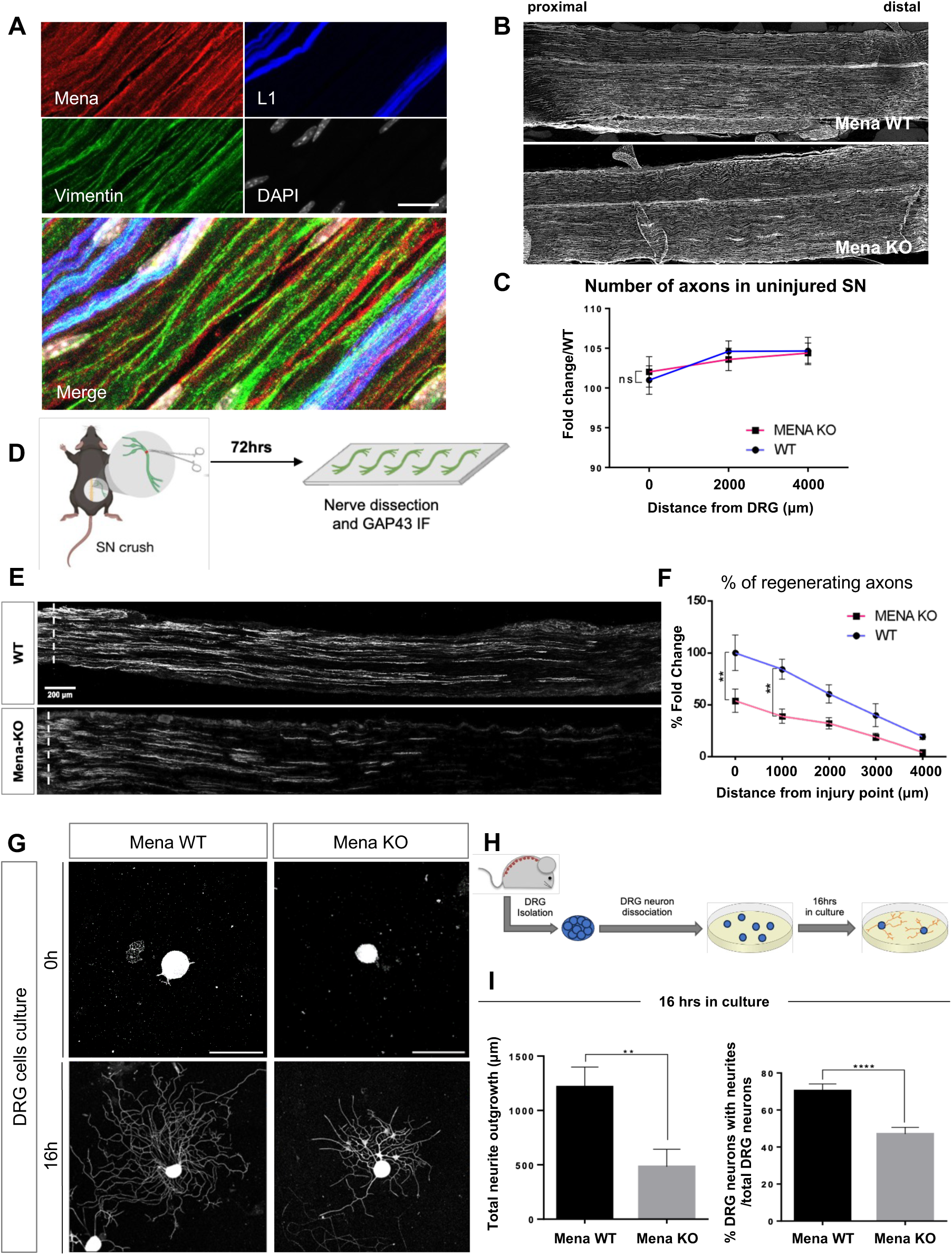
Mena- deficiency affects the regenerative capacity of adult SN axons after injury in vivo. **A.** IF and confocal microscopy on longitudinal cryosections from mouse SNs. Mena (red) is highly expressed in axons that are visualized by Vimentin (green) and L1 (blue). DAPI staining (grey) depicts the nuclei of glial cells within the nerve (Schwann cells). Scalebar: 50μm. **B.** IF and confocal microscopy of Neurofilament on longitudinal cryosections from mouse SNs, noting the proximal and distal site from the respective DRG (Dorsal Root Ganglion). Scalebar: 1000μm. **C.** Quantification of Neurofilament positive axons of uninjured SNs, at different distance from the DRG, normalized to WT axons (n=4 biologically independent animals per group, two-way analysis of variance (ANOVA) with Sidak post hoc test; mean ± stDEV). **D.** Schematic representation of the *in vivo* SNC, which was followed by 3 days of regeneration (72hrs). 2MO animals were used. **E.** Micrographs showing representative longitudinal sections of WT and Mena KO SNs 72hrs after SNC, immunostained for GAP43. The dashed lines indicate the crush site. Scalebar: 200μm. **F.** Quantification of the percentage of axons past the crush site, normalized to the number of axons at the crush site plotted as a function of the distance from the crush site (Regeneration index) (n=4 biologically independent animals per group with a bilateral SNC; **p< 0.01, two-way analysis of variance (ANOVA) with Tukey’s post hoc test examined over three independent experiments; mean ± stDEV). **G**. *In vitro* culture of dissociated DRG neurons from 2MO WT and Mena KO mice, 4hrs and 16hrs after plating, with representative morphologies within each genotype (WT panels: i-iii and KO panels: iv-vi). Scalebar: 50μm. **H**. Schematic representation of the *in vitro* cultures of dissociated DRG neurons. **I.** Quantification of total neurite outgrowth in dissociated neurons (in μm) after 16hrs in culture (n=3 biologically independent cultures, where 375 WT neurons and 338 Mena KO neurons were assessed). **p< 0.01, Unpaired t-Test between Wild Type and Mena KO DRG neurons; mean ± stDEV, and quantification of cells with axons, over the total cell population after 16hrs *in vitro* (n=3 biologically independent cultures, where 375 WT neurons and 338 Mena KO neurons were assessed). ****p< 0.0001, ANOVA with Tukey’s post hoc test; mean ± stDEV.

In order to investigate whether Mena is important for regeneration of SN axons after injury, we employed a loss-of-function strategy, by generating mice with CRISPR/Cas9-mediated genetic ablation of Mena (Mena KO mice; Fig. S1B, C, D, E, F, Table S1). Similar to other Mena KO mice(Kwiatkowski et al. 2007; Lanier et al. 1999; Menzies 2004), these animals are viable and fertile and an initial characterization and comparison of their nervous system with that of WT littermates, revealed no gross morphological differences and defects (data not shown). Importantly, adult (2MO) Mena KO mice do not show differences in their SN axons under physiological conditions *in vivo* (Fig. 1B, C). We then used 2MO Mena KO and WT mice to examine the regenerative capacity of their axons after sciatic nerve crush *in vivo* (SNC). Regeneration was assessed using IF for GAP-43, and confocal microscopy on longitudinal nerve cryosections that were isolated from mice 72hrs after SNC(Serger et al. 2022) (Fig. 1D). We found that the number of Mena-deficient axons extending away from the crush site was significantly lower, compared to axons of WT animals (Fig. 1E), as indicated by the regeneration index, which was calculated by measuring the average GAP43 intensity in bins across at least 4 mm distal to the crush site, as previously described(Serger et al. 2022) (Fig. 1F). This observation denotes a reduced axonal capacity for growth and regeneration after injury of adult SNs in the absence of Mena.

In order to examine whether our observation is due to intrinsic or extrinsic factors in the absence of Mena, we isolated DRGs (Dorsal Root Ganglia) from 2MO Mena KO and WT mice, and plated dissociated neurons in culture, thus diminishing any potential influence from their microenvironment (Fig. 1H). Cytosine Arabinose (Ara-C) treatment was used to minimize glial survival and proliferation in neuronal cultures. The intrinsic ability of cells to re-grow their axons after isolation (axotomy) was assessed using IF for the axonal marker Tuj1 (β3-tubulin) and confocal microscopy (Fig. 1G). We found that approximately 75% of WT DRG neurons were able to grow axons 16hrs after plating, as opposed to 45% of their Mena KO counterparts, while the total neurite outgrowth was also significantly lower in Mena KO neurons (Fig. 1I). This result highlights the significance of Mena with respect to the intrinsic capacity of adult axons to grow and regenerate after injury.

### Mena interactions in adult SNs reveal the implication of the protein in axonal LT

To understand the molecular function of Mena in adult axons, and how it might be implicated in regeneration after injury, we next wanted to identify protein-protein interactions of Mena in adult SNs. However, the adult SN is very highly enriched in myelinating glial cells (Schwann cells; DAPI staining in Fig. 1A) that also express Mena. To examine the function of Mena specifically in adult axons, and independently of glia, we developed a protocol for axoplasm purification from SNs(Perlson et al. 2004; Rishal et al. 2010), diminishing contamination from glial cells (Fig. S2A, B). Using axoplasm from mouse SNs, purified from 2MO WT animals, we performed mass spectrometry after immunoprecipitation (IP) of Mena (Mena-IP) (Fig. 2A). Respective axoplasm from Mena KO animals was used as a negative Control IP (Fig. 2A). We identified 271 proteins significantly associated with Mena in adult axons (Fig. 2B; −(logP) value>1.7; dif>0.5) (Table S2). Gene Set Enrichment Analysis (GSEA, Biological Process) that was performed on gene lists generated by group comparisons(Yu et al. 2012), revealed the most significant functional categories of Mena-interacting proteins, among which mRNA translation and actin filament organization are found among the top positions (Fig. 2C). Further analysis of all significant Mena-interactors was performed using STRING-based clustering of the respective molecules(Van Dongen 2008), and visualization based on the Cytoscape software for biomolecular interaction networks(Shannon et al. 2003). All actin-regulating proteins (79 proteins, orange dots in Fig. 2B) clustered together (Cut-off 0.55; Inflation 2.5), and among those we identified numerous known interactors of Mena, including Actin, VASP, Profilin2(Vidaki et al. 2017; Barzik et al. 2005), as well as novel ones (Fig. S2C).

**Figure 2:**
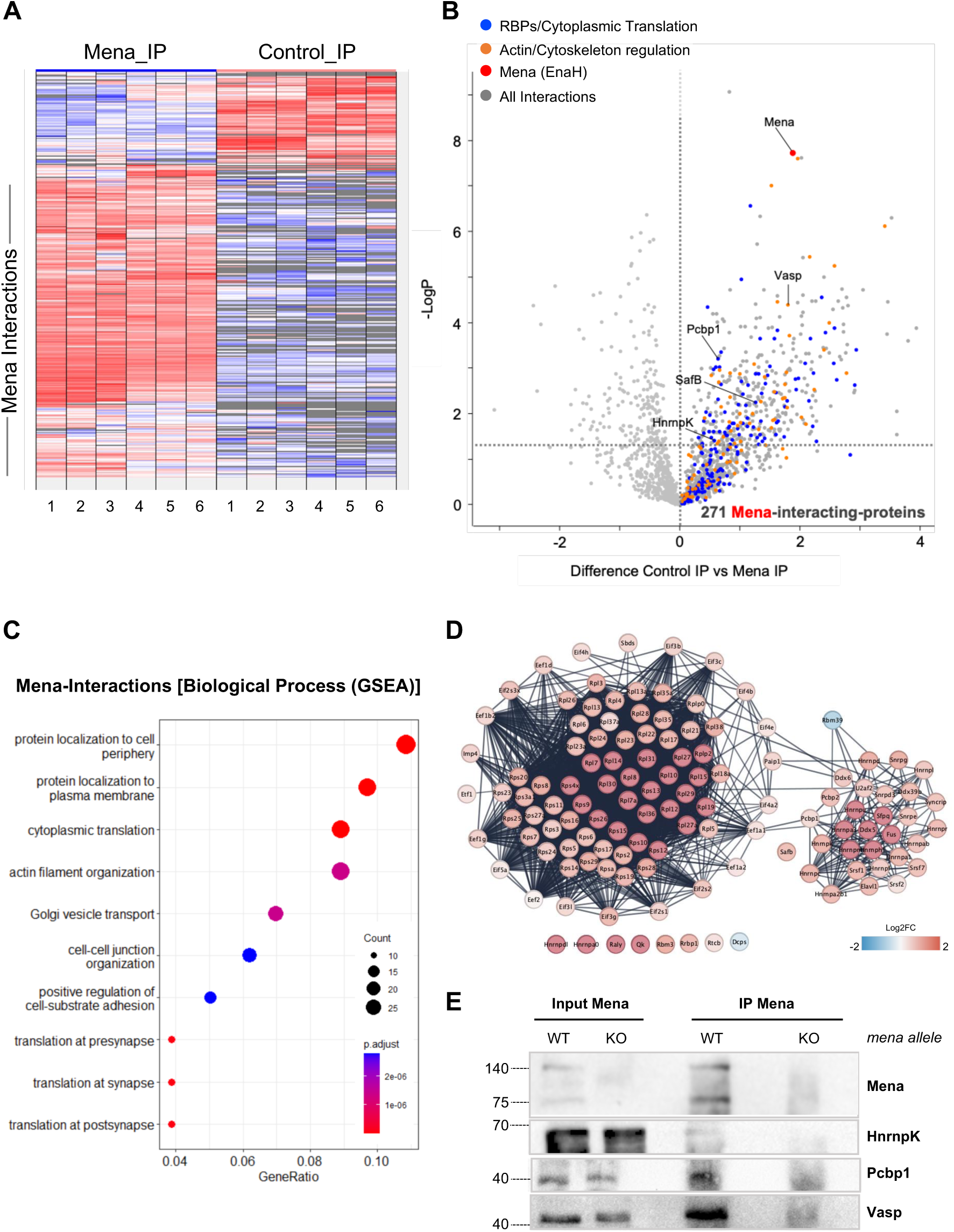
Mena interactions in adult SNs reveal the implication of the protein in axonal LT. **A.** Heat Map of proteins significantly interacting with Mena in adult SN axoplasms. (n=3 independent biological replicates and 2 technical repeats per sample). **B.** Volcano plot of proteins found in association with Mena vs Control IP (Mena KO axoplasm). 271 proteins were identified as significantly interacting with Mena in adult axons (-(logP) value>1.7; dif>0.5). Interactions with RBPs are highlighted in blue, including known interactions like HnrnpK, PCBP1, SafB. Interactions with cytoskeleton associated proteins are highlighted in orange (e.g. VASP). **C.** Gene Set Enrichment Analysis (GSEA, Biological Process) on gene lists generated by group comparisons. mRNA translation and actin filament organization are among the top categories of interactors. **D.** mRNA processing-related proteins interacting with Mena (124 proteins, blue dots in 2B) clustered in two major groups (Cut-off 0.85; Inflation 2.5), one with RBPs that are implicated in splicing, mRNA transport and translation regulation (e.g. Hnrnp proteins and all the known components of the Mena-RNP, like HnrnpK, Pcbp1, SafB, HnrnpA2B1, HnrnpM, etc), and a second including ribosomal proteins of the small and large subunits, and translation initiation and elongation factors (e.g. EiF4 family members, eEF2, etc). **E.** coIP experiment that reveals the interaction between Mena with HnrnpK and Pcbp1. VASP was used as a positive control for the IP efficiency.

On the other hand, all Mena-interacting proteins that are associated to mRNA processing (124 proteins, blue dots in Fig. 2B) clustered in two major groups (Cut-off 0.85; Inflation 2.5), one including RBPs that are implicated in splicing, mRNA transport and translation regulation (e.g. Hnrnp proteins and all the known components of the Mena-RNP, including HnrnpK, Pcbp1, SafB, HnrnpA2B1, HnrnpM, and others(Vidaki et al. 2017)), and a second one that includes ribosomal subunits and translation initiation and elongation factors (e.g. EiF4 family members, eEF2, etc) (Fig. 2D). Using IF and confocal microscopy, as well as axoplasm purifications and Mena co-immunoprecipitation (coIP) assays, we validated the association of Mena with several RBPs, including known components of the developing Mena-RNP, like HnrnpK and Pcbp1, in adult SN axons (Fig. 2E, Fig. S2D, E).

Overall, our findings uncover multiple novel interactions for Mena in adult axons, as well as numerous previously established ones, suggesting the implication of the protein in adult axon translation regulation.

### The absence of Mena disrupts mTOR LT and downstream signaling, as an acute response to injury

The significant association of Mena with proteins that are implicated in multiple aspects of RNA metabolism, including translation and ribosomal structure, suggests a potential effect of the protein in LT in adult axons. Therefore, we examined the role of Mena in axonal protein synthesis after injury. We employed an *ex vivo* injury model, in which SNs are cut into smaller segments-after dissection and removal of the respective DRGs, and subsequently incubated in culture medium for up to 24hrs(Terenzio et al. 2018; Perlson et al. 2005) (Fig. 3A). This method allowed us to study the intrinsic capacity of axons for local protein synthesis by monitoring even minor molecular changes that occur rapidly in axons following injury, independently of the cell body and the complex extracellular microenvironment. Using this system, followed by axoplasm isolation, we visualized axonal responses to injury at selected timepoints (0hrs, 2hrs, 4hrs and 24hrs) (Fig. S3A, B, C and data not shown). The acute injury response of SN segments was optimally observed using western blot analysis at 4hrs post injury, by the elevation of specific proteins that are known to be locally translated after injury, including Vimentin, Importin-β1, and mTOR(Perlson et al. 2004; Terenzio et al. 2018), as well as by the increased phosphorylation of ribosomal protein S6, which demarcates active protein synthesis(Terenzio et al. 2018) (Fig. S3A). Both Vimentin, and Importin-β1 have been shown to be locally synthesized in axons of the PNS after axotomy, in order to bind phospho-ERK (extracellular-regulated kinase, ERK) and retrogradely transport it to the nucleus, where it activates regeneration-associated genes (RAGs)(Perlson et al. 2004; Smith et al. 2020; Perlson et al. 2005). Concomitantly, mTOR (mammalian Target of Rapamycin), which is a major regulator of protein homeostasis, has also been shown to be locally translated after axotomy in the PNS, to modulate local protein synthesis and facilitate regeneration(Terenzio et al. 2018). To confirm that the observed protein level changes in our *ex vivo* injury model, were indeed due to LT events, we used anisomycin to block translation (Fig. S3B, C and data not shown).

**Figure 3:**
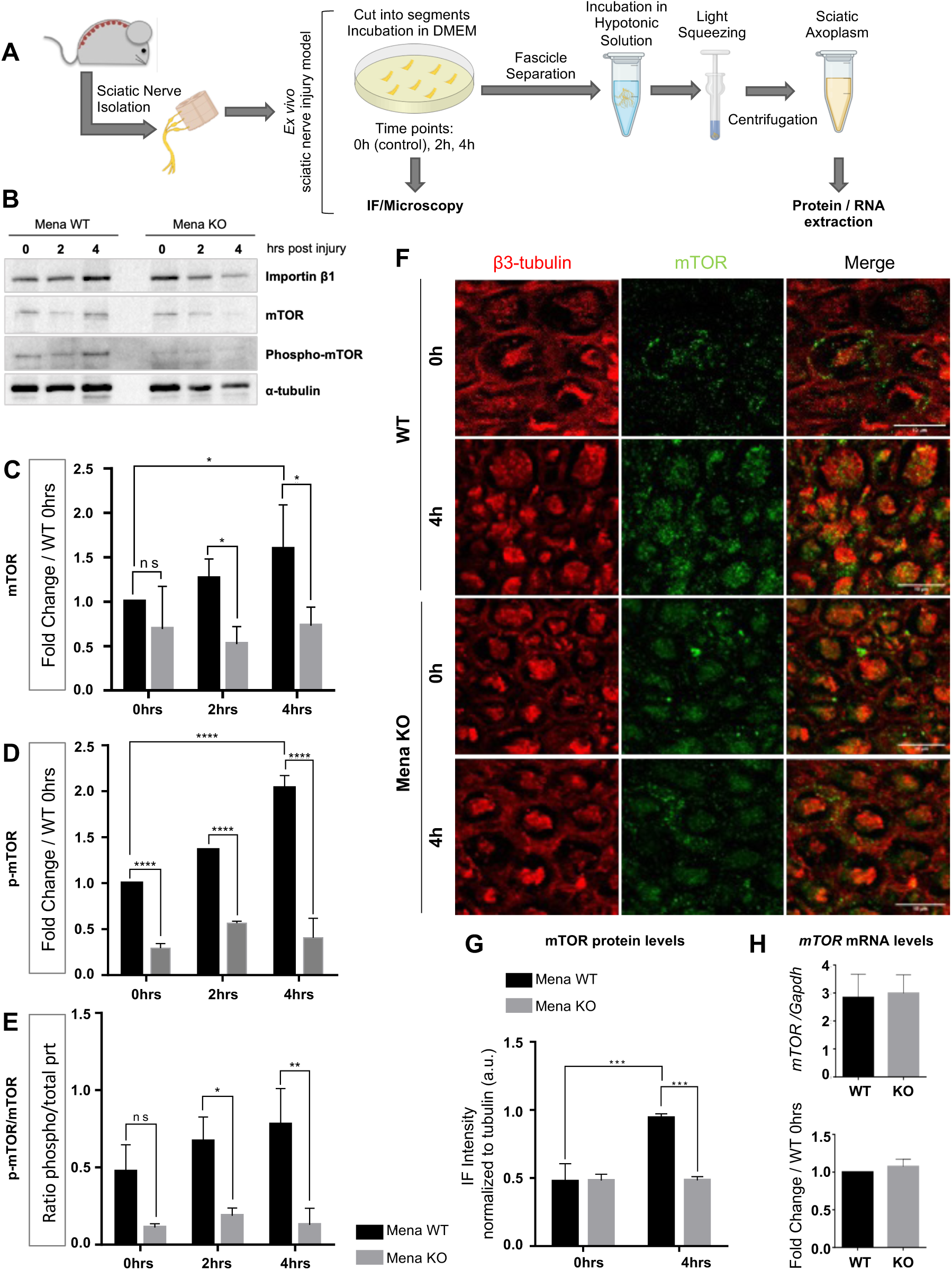
Mena-ablation disrupts mTOR local translation, as an acute response to injury ex vivo. **A.** Schematic representation of the *ex vivo* injury model used in the study. At 0hrs in culture the axons were considered as naïve, and the optimal acute response to injury was observed 4hrs after segmentation of SNs. **B.** Western blot analysis of mTOR and Importin-β1 proteins after *ex vivo* injury of axons from WT and Mena KO SNs from 2MO mice. **C.** Local translation of mTOR, as well as mTOR phosphorylation (Ser2448) (**D**) and the ratio of p-mTOR/total-mTOR (**E**) are significantly decreased in axons after injury, due to Mena-ablation (n=4 biologically independent replicates per condition. *p < 0.05; **p < 0.01, ****p < 0.0001, ANOVA with Tukey’s post hoc; mean ± stDEV). **F.** IF of mTOR (green) protein levels and distribution in cross sections from WT and Mena KO SN segments pre- and post-injury *ex vivo*, reveals significant reduction of mTOR in the absence of Mena (quantifications in Figure S3). Scalebar: 10μm. **G.** Quantification of the IF signal of mTOR, normalized to tubulin intensity of fluorescence at 0hrs and 4hrs post-injury *ex vivo*. (n=3 biologically independent replicates per assay, per condition. ***p<0.001, ANOVA with Tukey’s post hoc; mean ± stDEV). **H.** Quantification of *mTOR* mRNA in Mena KO axons (normalized to total *Gapdh* mRNA and to WT *mTOR* mRNA) reveals no alteration in the mRNA levels (n=3 biologically independent replicates per assay, per condition, ANOVA with Mann-Whitney; mean ± stDEV).

Next, we used this system to characterize the acute response of Mena KO axons to *ex vivo* injury. Western blot analysis revealed that Mena-ablation significantly affects LT of Importin-β1 (Fig. 3B, Fig. S3D), as well as mTOR (Fig. 3B, C) 4hrs post injury. The fact that both proteins are not affected by the absence of Mena in naive axons (Fig. 3B, C, Fig. S3D, 0hrs), supports the hypothesis that Mena is important for LT events in response to injury. In agreement with the lower levels of mTOR 4hrs post-injury, phosphorylation of mTOR (p-mTOR; Ser2448) was also affected, leading to significantly decreased levels of active mTOR in Mena KO axons (ratio of p-mTOR/mTOR) (Fig. 3D, E). The inability of Mena KO axons to elevate their mTOR protein levels in response to injury was further confirmed by IF experiments and confocal microscopy, on cryosections from SN segments after 4 hrs of *ex vivo* injury (Fig. 3F, G and Fig. S3E). Using Pearson’s Coefficient analysis of colocalization between mTOR and axonal β3-Tubulin IF signal, we confirmed that the observed increase in IF signal was indeed axonal mTOR (Fig. S3F, G). In order to verify that mTOR protein levels post injury are affected locally as a result of defective translation and not transcript abundance, we performed quantitative PCR (qPCR) to monitor *mTOR* mRNA levels in WT and Mena KO SN axoplasm. We observed that *mTOR* mRNA levels were not affected by the absence of Mena in axons (Fig. 3H). This suggests that transcription, transport and axonal localization of *mTOR* transcripts was not perturbed by Mena-ablation, in contrast to their impaired LT upon injury.

Given the fact that mTOR is a central regulator of cell growth, subcellular protein synthesis, and regeneration(Saxton and Sabatini 2017; Terenzio et al. 2018; Altas et al. 2022), we aimed to examine further key components of the mTOR pathway with respect to their levels and changes in axons pre-and post-injury, and potential alterations caused by Mena-ablation. For example, mTOR regulates the phosphorylation of the 70 kDa protein kinase (p70 S6k, S6K1), which acts on ribosomal protein S6 and is involved in controlling the translation of multiple mammalian mRNAs that encode ribosomal proteins(Al-Ali et al. 2017; Wang et al. 2001). Using western blot analysis after *ex vivo* injury of SNs from 2MO WT and Mena KO animals, we found considerably deregulated S6K1 due to Mena ablation (Fig. 4A, B). More specifically, Mena KO axons display significantly lower levels of S6K1 4hrs post *ex vivo* injury, which was also verified by IF experiments and confocal microscopy, on cryosections from SN segments after 4 hrs of *ex vivo* injury (Fig. S4A, B, C). This decrease is the result of limited LT of S6K1 in Mena KO axons, as indicated by puromycin pulse-chase experiments. Briefly, low concentration of puromycin was added to the medium of cut SN segments and its incorporation was subsequently traced 4hrs after injury, using IF and confocal microscopy on cross, or longitudinal sections of SN segments (Fig. 5A). A time point of 0hrs, as well as 4hrs without puromycin addition to the medium, or with puromycin and anisomycin to block translation were used as internal controls. This assay allowed us to detect S6K1 signal and colocalization with puromycin (newly synthesized S6K1) and axonal β3-Tubulin (Pearson’s Coefficient) (Fig. S4A, B, C, D).

**Figure 4:**
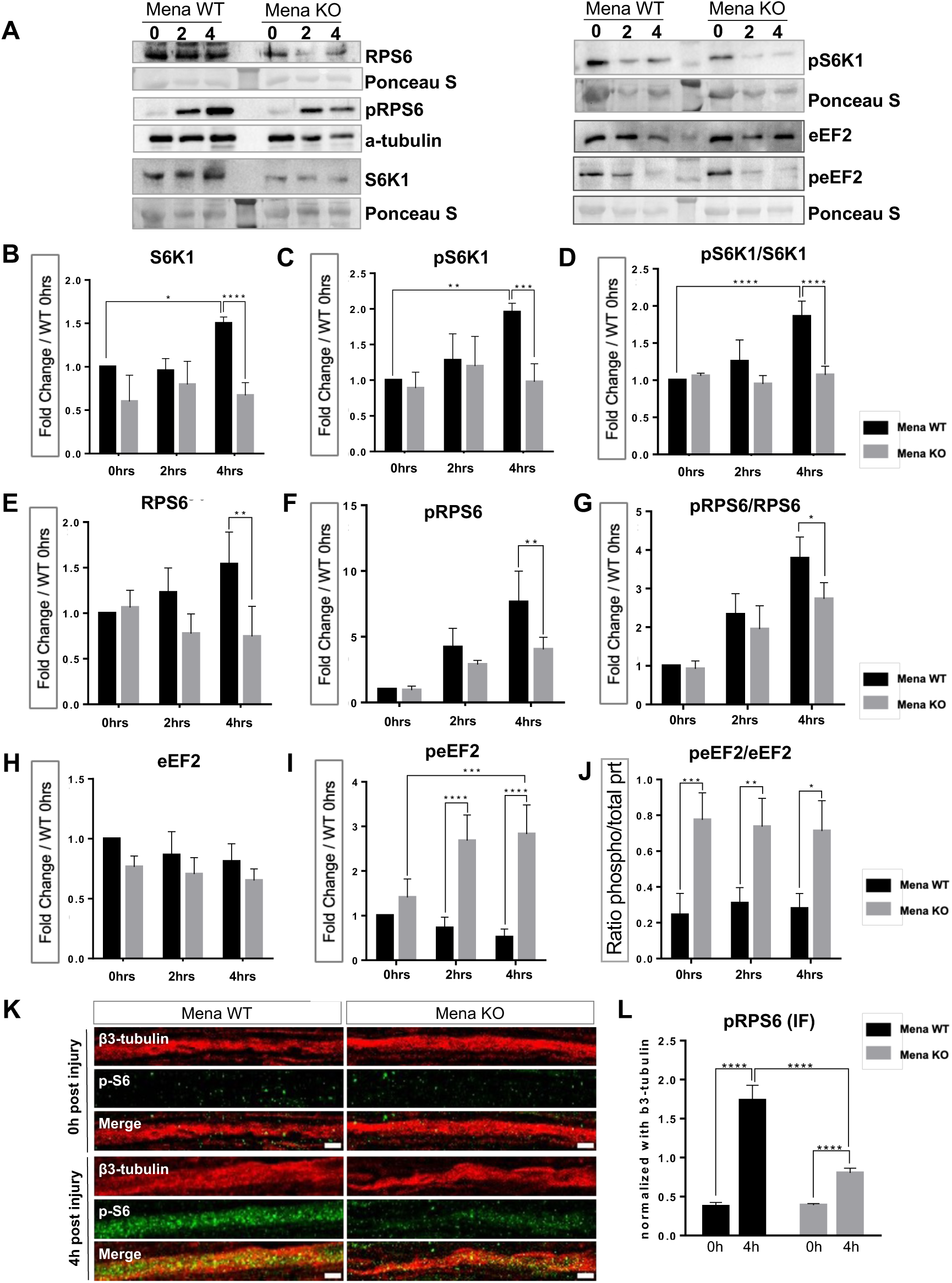
The absence of Mena deregulates mTOR downstream signaling after injury *ex vivo*. **A.** Western blot analysis of mTOR effector proteins S6K1 (p70-S6 Kinase), ribosomal protein S6, eEF2, eEF2 kinase, and their phosphorylated forms (p-S6K1; Ser371, p-S6; Ser240/244 and p-eEF2; Thr56 respectively). Quantification of the protein levels reveals that S6K1 levels do not increase as anticipated in Mena KO axons in response to injury, neither does phosphorylation (**B, C**) and activation of the kinase (**D**, p-S6K1/t-S6K1 ratio). **E.** S6 levels appear to be significantly decreased in Mena KO axons 4hrs post-injury, as do levels of pS6 (**F**) and the p-S6/t-S6 ratio (**G**) that represents S6 activity and demarcates active translation. **H.** Total levels of eEF2 are significantly downregulated in axons of Mena KO nerves, 4hrs post- injury, while phosphorylated-eEF2 (p-eEF2; Thr56), which is inactive, is significantly elevated in the absence of Mena (**I, J**). (n=4 biologically independent replicates per condition. *p < 0.05; **p < 0.01, ***p < 0.001, ****p < 0.0001, ANOVA with Tukey’s post hoc; mean ± stDEV). **K, L.** IF and confocal microscopy on longitudinal cryosections from SN axons at 0 and 4hrs post injury reveal the increased phosphorylation of S6 in WT but not Mena KO axons, further highlighting their limited translational activity (n=5 biologically independent replicates per condition. ****p < 0.0001 (ANOVA with Tukey’s post hoc; mean ± stDEV). Scalebar: 5μm

**Figure 5:**
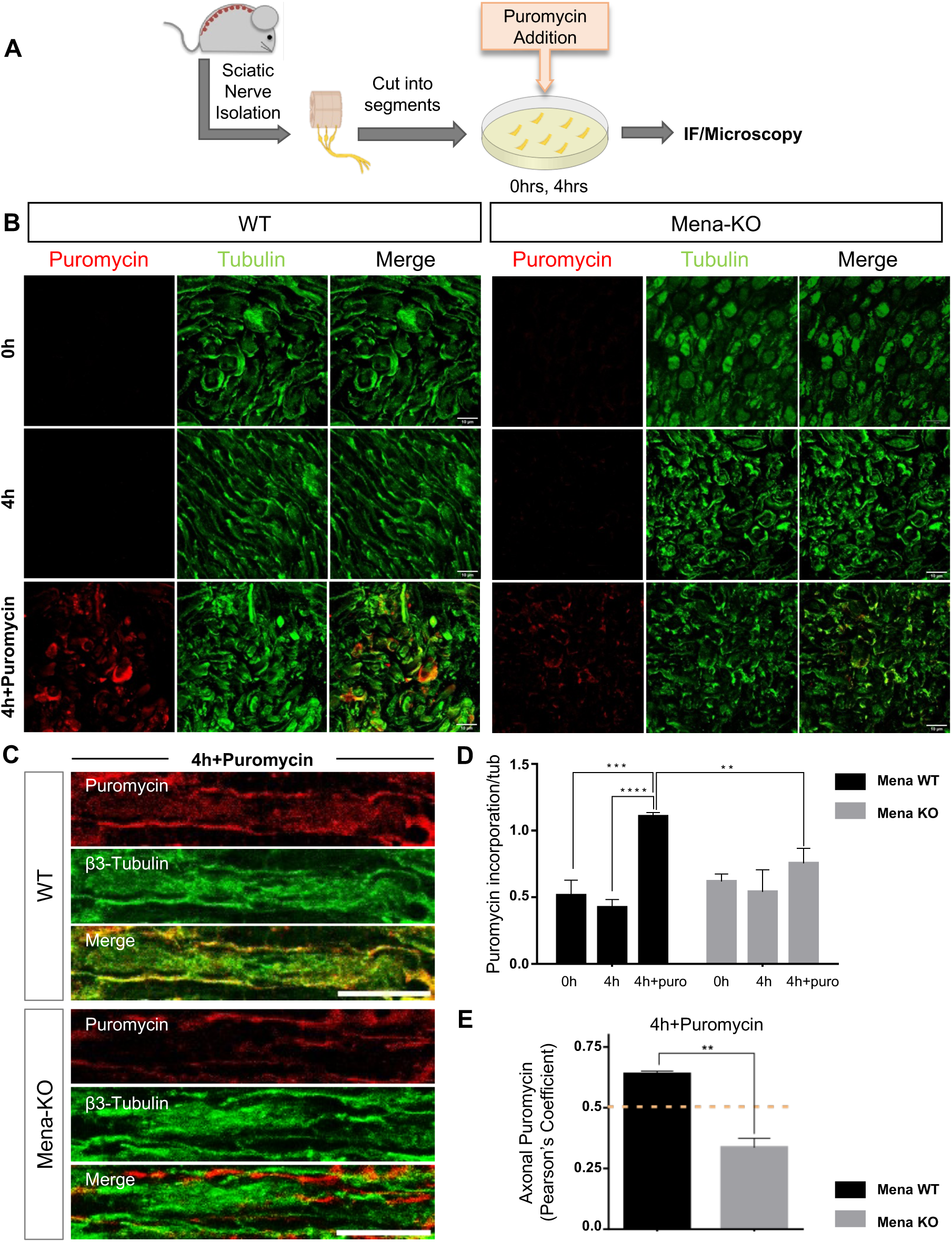
Mena-ablation results in decreased puromycin incorporation, confirming global protein synthesis defects in adult SN axons after injury. **A.** Schematic representation of the puromycin pulse-chase experiment that was performed in combination with *ex vivo* injury on SNs from WT and Mena KO mice. IF and confocal microscopy on transverse (**B**) or longitudinal sections (**C**) from sciatic nerves, was used to visualize puromycin (red) levels and distribution in axons, as a marker of novel protein synthesis in response to injury. Axons were additionally labeled with β3-Tubulin (green) to visualize axonal puromycin incorporation. **D**. Mena-deficient axons display significantly lower puromycin incorporation, compared to WT axons (Intensity values of puromycin were normalized to total β3-Tubulin values; n=3 biologically independent replicates per condition. **p < 0.01, ***p<0.001, ****p < 0.0001, ANOVA with Tukey’s post hoc; mean ± stDEV). Scalebar: B=10μm, C=50μm. **E**. Pearson’s Coefficient was assessed to verify colocalization between puromycin and axonal β3-Tubulin (n=3 biologically independent replicates per condition. **p < 0.01, ANOVA with Tukey’s post hoc; mean ± stDEV).

In good agreement with compromised mTOR activity the phosphorylated S6K1 (p-S6K1; Ser371) was also reduced in the absence of Mena (Fig. 4A, C, D). Similarly, ribosomal protein S6 levels were significantly decreased in Mena KO axons 4hrs after injury (Fig. 4A, E), and phosphorylation of S6 (p-S6; Ser240/244) that should be increased in response to injury, failed to do so (Fig. 4A, F). This shifted the overall activation of S6 (p-S6/t-S6 ratio, Fig. 4G) that demarcates active protein synthesis, towards significantly lower levels in the absence of Mena. This observation was further validated by IF experiments and confocal microscopy, on cryosections from SN segments after 4 hrs of *ex vivo* injury, where p-S6 signal appears to be significantly lower in Mena KO axons (Fig. 4K, L).

mTOR-mediated activation of S6K1, not only activates S6, but also inhibits eEF2 Kinase (eEF2K) function, thereby regulating eEF2 phosphorylation levels and promoting peptide chain elongation and protein synthesis(Smith et al. 2020; Yang et al. 2014). Our study revealed that Mena KO axons have significantly lower total eEF2 levels 4 hrs post-injury (Fig. 4A, H), and highly elevated levels of phosphorylated eEF2 (p-eEF2; Thr56) (Fig. 4A, I). This resulted in overall diminished eEF2 activity (p-eEF2/t-eEF2 ratio, Fig. 4A, J), despite the fact that total levels of eEF2K remain unaffected (Fig. 4A, Fig. S4E). Therefore, hyper-phosphorylated eEF2 in Mena KO axons may be the result of decreased mTOR levels and function, and thus inability to inactivate eEF2K.

Taken together, our data demonstrate a significant role for Mena in LT and activity of mTOR and its downstream effectors in response to injury of SN axons.

### Mena-deficiency results in diminished global protein synthesis in adult SN axons after injury

Our puromycin pulse-chase experiment to verify *de novo* synthesis of S6K1 in SN axons revealed decreased puromycin IF signal in Mena KO axons (Fig. S4A, B, C). Given the fact that the mTOR signaling pathway is a major regulator of protein synthesis under physiological and pathophysiological conditions, and a key intrinsic mechanism that modulates regeneration, we reasoned that the inability of Mena KO axons to regenerate after injury *in vivo*, may be essentially attributed to defective global translation. So next, we assessed the intrinsic ability of axons for protein synthesis after injury, and the implication of Mena in the process, by employing pulse-chase puromycin incorporation in newly synthesized peptides, after *ex vivo* injury of WT and Mena KO SNs. As described above, low concentration of puromycin was added to the medium of SN segments and its incorporation was subsequently traced 4hrs after injury, using IF and confocal microscopy on cross, or longitudinal sections of SN segments (Fig. 5A, B, C). A time point of 0hrs, as well as 4hrs without puromycin addition to the medium were used as internal controls. To verify that puromycin incorporation was quantified only in axons and not glial cells present in the SN segments, we analyzed the colocalization of puromycin signal with β3-Tubulin, using Pearson’s Coefficient Correlation. We found that Mena KO axons displayed significantly less puromycin incorporation, compared to WT axons (Fig. 5B, C, D, E), uncovering their intrinsic inability to synthesize proteins *in situ*.

Taken together, our findings support the notion that Mena is necessary for local regulation of the mTOR/S6K1 pathway and subsequent axonal protein synthesis in adult axons of the PNS in response to injury.

### Mena-deficient axons of SNs display crucial alterations in their local proteome

In order to gain further mechanistic insight into Mena function with respect to local regulation of the mTOR/S6K1 pathway and downstream signaling, we next wanted to examine the axonal proteome of SN axons pre- and post-injury *ex vivo*, so as to detect potential changes due to Mena ablation. Using axoplasm preparations from 2MO WT and Mena KO animals, and quantitative Mass Spectrometry we identified numerous proteins differentially expressed in WT vs Mena KO axons (Table S3). More specifically, we found 210 proteins enriched in the axoplasm of naïve WT axons, as opposed to 43 proteins enriched in the axoplasm of naïve Mena KO axons (Fig. 6A; −(logP) value>1.7; dif>0.5). After *ex vivo* injury 1214 proteins were enriched in the axoplasm of WT axons, as opposed to 432 proteins enriched in the axoplasm of Mena KO SNs (Fig. 6B; −(logP) value>1.7; dif>0.5). Gene Set Enrichment Analysis (GSEA, Biological Process) that was performed on the differentially expressed (DE) gene lists generated by group comparisons, revealed the most significant functional categories of the enriched proteins in each condition (Fig. S5A, B). Interestingly, we observed that numerous proteins of the PI3K/Akt/mTOR pathway, were DE in WT and Mena KO axons post-injury, but not in naïve axons (Fig. 6A, B and Fig. S5B), potentially as a result of the decreased ability of Mena KO SNs to regulate LT as an acute injury response. More specifically, in good agreement with our findings, mTOR, and downstream effectors of mTOR, like subunits of S6K1 (Rps6ka1, Rps6ka2, Rps6kb1, Rps6kb3), the ribosomal protein S6 (RpS6), and translation elongation factor eEF2, are also DE in WT vs Mena KO axons 4hrs post-injury (Fig. 6B and Fig. S5B). Concomitantly, we found that members of the Akt serine-threonine protein kinase family (Akt1, Akt3) are elevated in WT axons 4hrs post-injury, contrary to the respective Mena KO axons, where no such response was observed (Fig. 6B and Fig. S5B). Further validation of our Mass Spectrometry data by western blot analysis after *ex vivo* injury of adult SNs, revealed that indeed, Akt levels are slightly decreased in axons of Mena KO animals 4hrs after injury, while phosphorylation of Akt (p-Akt; Ser473), and thus activity of the protein (p-Akt/t-Akt ratio) were also significantly lower in the absence of Mena (Fig. 6C, E, F, G).

**Figure 6:**
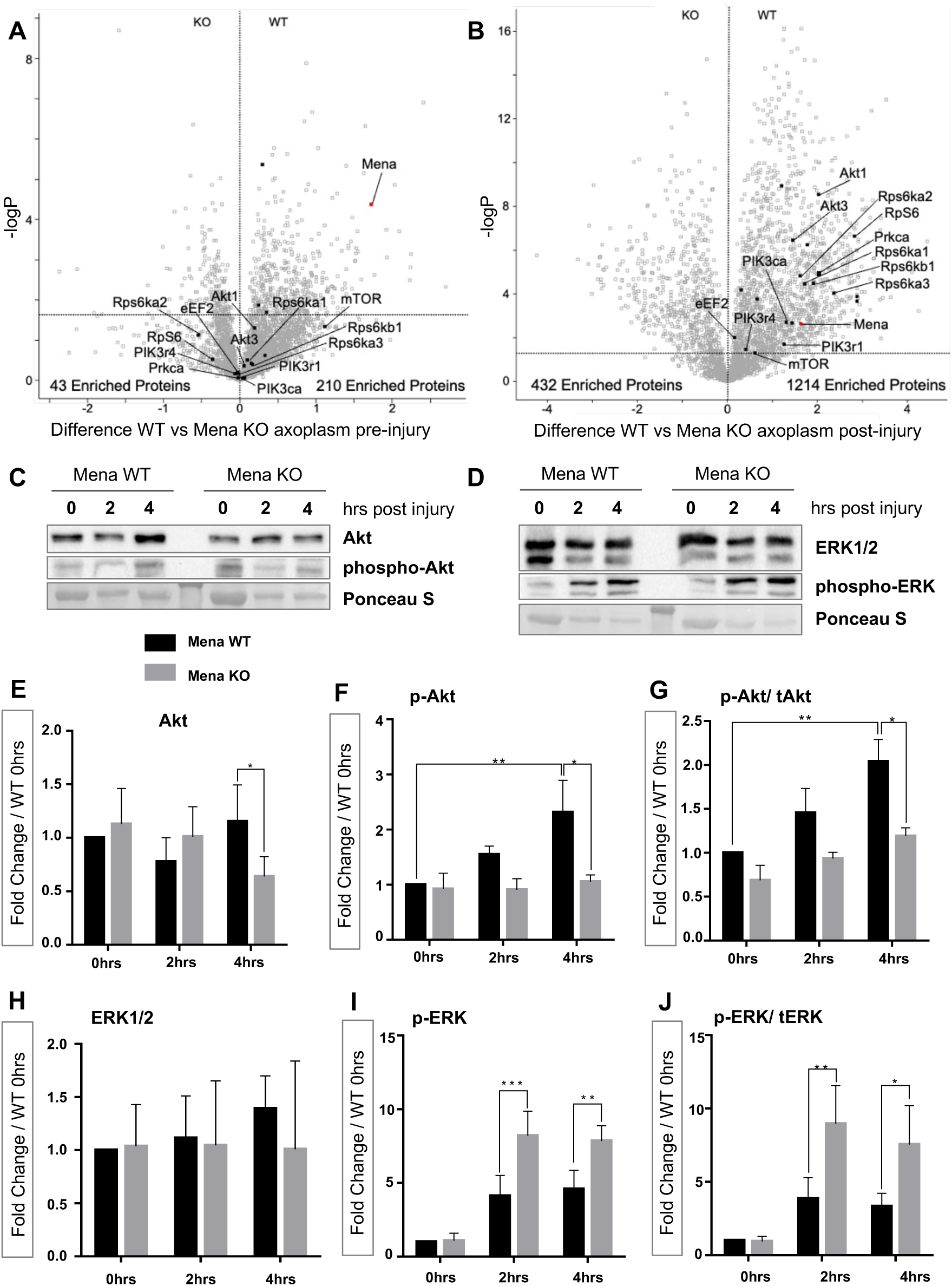
Significant alterations in the local proteome of SN axons in the absence of Mena. **A.** Volcano plot of differentially expressed (DE) proteins in SN axons of 2MO WT and Mena KO mice pre-injury. Members of the mTOR pathway that grossly regulate translation and regeneration are highlighted, and are not significantly different due to Mena-ablation. **B.** Volcano plot of differentially expressed proteins in SN axons of 2MO WT and Mena KO mice 4hrs post-injury. Members of the mTOR pathway are highlighted and appear to be DE between WT and Mena KO axons 4hrs post-injury. **C.** Western blot analysis of total and phosphorylated Akt (p-Akt; Ser473) levels, as well as total and phosphorylated ERK (p-ERK; Thr202/Tyr204) levels (**D**) in response to injury. Total Akt is significantly downregulated 4hrs after injury in Mena KO axons (**E**), as is its phosphorylation status (**F**) and overall activity (**G**). Total ERK (ERK1/2) levels are not affected by Mena-ablation (**H**), but ERK phosphorylation (**I**) and the ratio of p-ERK/t-ERK that mirrors its activity (**J**), are significantly elevated in Mena KO axons post-injury. (n=4 biologically independent replicates per condition. *p < 0.05; **p < 0.01, ANOVA with Tukey’s post hoc; mean ± stDEV)

Another signaling pathway that has been shown to be crucial for regeneration after injury, activated independently of mTOR, is that of extracellular signal-regulated kinase (ERK). Activation of ERK is required for axotomy-induced growth cone reformation after lesion, and its inhibition impairs axon regeneration in vivo(Hausott and Klimaschewski 2019). Moreover, ERK regulates phosphorylation of p90-S6 kinase (p90 RSK1) that can further phosphorylate and inhibit eEF2K, promoting translation independently of mTOR(Wang et al. 2001). We found that ERK protein levels (ERK1/2) remain unaffected by Mena-ablation in SN axons pre- and post-injury *ex vivo* (Fig. 6D, H). However, ERK phosphorylation (Thr202/Tyr204) is significantly upregulated in Mena KO axons post injury (Fig. 6D, I), indicating increased ERK activity due to the absence of Mena (Fig. 6J). This may be mediated by decreased levels of PKA in Mena KO axoplasms, as indicated by the differential expression of the kinase in our Mass Spectrometry data (Prkca levels elevated in WT, but not Mena KO axons post injury; Fig. 6A, B and Fig. S5B). This could be a potential compensatory mechanism that axons activate in the absence of Mena, to sustain their homeostasis upon trauma.

Taken together, our findings indicate that Mena ablation affects the Akt/mTOR signaling axis, leading to decreased LT and regeneration after injury; despite increased activation of ERK, Mena KO axons are unable to compensate for the deficiency.

### Mena associates with *Pi3kr1* mRNA and regulates LT of PI3K-p85 in response to injury of SN axons

Interestingly, our Mass Spectrometry data revealed that several subunits of PI3K (phosphoinositide 3-kinase) were DE in Mena WT versus KO axoplasms in response to injury (Fig. 6A, B and Fig. S5B). Given the fact that Akt activation (Ser473-phosphorylation) and subsequent mTOR signaling are PI3K-dependent, we reasoned that PI3K may be regulated by Mena, thus explaining the defective Akt/mTOR signaling and subsequent translation regulation. PI3K is an obligate heterodimer with a regulatory subunit (p85) and a catalytic subunit (p110). The primary function of p85 is to bind, and stabilize the p110 catalytic subunit, regulating its activity until required(Sánchez-Alegría et al. 2018; Taniguchi et al. 2010). Since the DE proteins in our dataset primarily belong to the regulatory subunit of PI3K (Pik3r1, Pik3r4; Fig. 6A, B), we tested the expression levels of PI3K-p85 (Pik3r1) in axons from WT and Mena KO SNs, pre- and post-injury *ex vivo*. We found that PI3K-p85 is locally translated in axons 4hrs post-injury, as indicated by the elevated levels of the protein that are diminished when translation is blocked with anisomycin (Fig. 7A, B). This *de novo* translation was not observed in Mena KO axons, as demonstrated by western blot and IF analysis (Fig. 7A, B, C, D).

**Figure 7:**
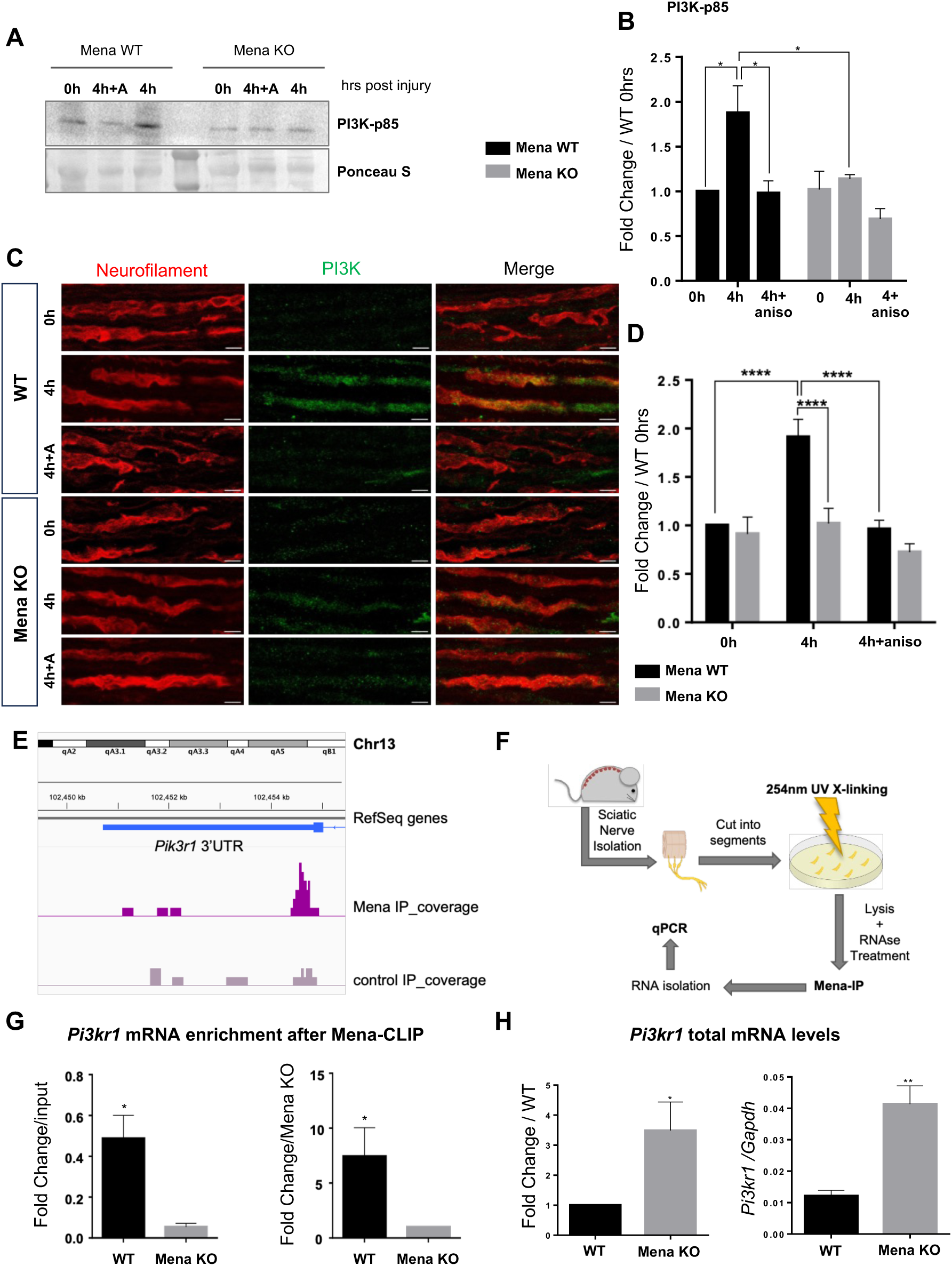
Mena associates with *Pik3r1* mRNA and regulates its LT in response to injury. **A.** Western blot analysis of PI3K-p85 subunit levels in WT and Mena KO axons of SNs, pre- and post-injury. **B.** Quantification of PI3K-p85 pre- and 4hrs post-injury with the addition of Anisomycin to block translation, reveals that the protein is locally translated in axons in response to injury *ex vivo*, but fails to do so in the absence of Mena. (n=4 biologically independent replicates per condition. *p < 0.05; **p < 0.01, ANOVA with Tukey’s post hoc; mean ± stDEV). **C.** IF and confocal microscopy of PI3K-p85 subunit (green) and Neurofilament (red) on longitudinal cryosections from WT and Mena KO SNs, at 0hrs and 4 hrs post injury, with and without Anisomycin addition. Scalebar: 5μm. **D.** Quantification of IF signal intensity of PI3K-p85, normalized to total Neurofilamnet intensity levels and over WT intensity at 0hrs reveals significant protein levels increase, as a result of LT in WT but not Mena KO axons 4hrs post-injury. (n=4 biologically independent replicates per condition. ****p < 0.0001 (ANOVA with Tukey’s post hoc; mean ± stDEV). **E.** Mena-HITS CLIP RNA sequencing peaks in the 3’UTR of *Pik3r1*, indicate a regulatory role of the interaction between Mena and the mRNA of the PI3K-p85 subunit. **F.** Schematic representation of the Mena-CLIP-qPCR experiment performed on SN axons. **G.** Enrichment of the *Pik3r1* transcript in the Mena- CLIP fraction over input and over the Mena KO control axons. (n=4 biologically independent replicates per condition. *p < 0.05; ANOVA with Mann-Whitney; mean ± stDEV. **H.** Total *Pik3r1* transcript levels are significantly elevated in axons of Mena KO SNs, compared to WT (n=4 biologically independent replicates per condition. *p < 0.05; **p < 0.01 (ANOVA Mann-Whitney; mean ± stDEV)

A previous study on the mRNAs that are regulated by Mena in developing axons, identified the *Pik3r1* mRNA of the regulatory PI3K-p85 subunit as being Mena-RNP-associated (Mena- HITS CLIP RNA Sequencing)(Vidaki et al. 2017). Further analysis of the particular set of data revealed numerous significant binding peaks of the Mena-RNP in the 3’UTR of the *Pik3r1* transcript (Fig. 7E), in good agreement with previous data supporting that Mena regulates translation of specific mRNAs, via the formation of a RNP that includes translation regulators, such as HnrnpK and PCBP1(Vidaki et al. 2017). Given the fact that the specific interactions of Mena are conserved in adult SN axons, as demonstrated by Mena-coIP Mass Spec and respective validating experiments in this study (Fig. 2B, C, D, E and Fig. S2D, E), we reasoned that Mena may regulate translation of the *Pik3r1* mRNA in adult axons. Using axoplasm preparations and Mena-IP after UV Crosslinking (Mena-CLIP) of SNs (Fig. 7F), we employed quantitative RT-PCR and found that the *Pik3r1* mRNA is indeed in complex with Mena in adult SN axons (Fig. 7G, CLIP/input and CLIP/KO).

To determine the abundance of the *Pik3r1* transcript in the axoplasm of adult Mena KO SNs, we isolated total RNA and performed quantitative RT-PCR. We observed that the total levels of the *Pik3r1* mRNA are significantly higher in Mena KO axons, compared to the respective WT axons (Fig. 7H). The increased abundance of the mRNA in the absence of Mena likely arises as a consequence of elevated transcription, increased mRNA stability, or both, potentially triggered by impaired translation, and suggests that Mena ablation does not affect axonal transport and localization of the transcript. Our findings indicate that the decreased PI3K-p85 protein levels observed after injury in Mena KO axons, are not due to low mRNA levels and/or defective mRNA transport, but rather due to the inability of those axons to translate the *Pik3r1* transcript locally, leading to its accumulation in the axoplasm.

Collectively, our data support the notion that Mena is necessary for LT of PI3K-p85 and local regulation of the PI3K/AKT/mTOR/S6K1 pathway, affecting the intrinsic capacity of adult PNS axons for protein synthesis and regeneration (Fig. S6).

## Discussion

The ability of axons to synthesize proteins locally on demand has been inextricably related to their intrinsic capacity for growth and regeneration after injury(Verma et al. 2005; Jung et al. 2012; Triantopoulou and Vidaki 2022; Nagano and Araki 2021). However, the molecular components and the mechanisms that modulate LT in adult axons remain only poorly understood. Our study uncovers that Mena, a member of the Ena/VASP family of proteins(Krause et al. 2003; Drees and Gertler 2008), directs LT of the PI3K-p85 subunit and activation of the PI3K/mTOR pathway in SN axons as an acute response to injury. We show that this function is pivotal for further axonal LT and regeneration, as highlighted by the incapacity of Mena-deficient SNs to synthesize proteins *de novo* and regenerate after injury. It has been previously shown that Mena has a dual function: it regulates cytoskeletal dynamics, via promoting actin filament polymerization (Dent et al. 2011; Kwiatkowski et al. 2007; McConnell et al. 2016; Menzies 2004), and local mRNA translation, via the formation of a RNP complex with RBPs and specific cytosolic mRNAs in developing axons(Vidaki et al. 2017). Both cellular processes are pivotal for axon regeneration(Smith et al. 2020; Akram et al. 2022; Bradke et al. 2012), thereby Mena could function as a key molecular coordinator of the two. However, the actin-regulating function can be partially compensated in neurons by other members of the Ena/VASP family, as previously described(Kwiatkowski et al. 2007; Lebrand et al. 2004; Menzies 2004), whereas the ability to associate with RBPs and regulate mRNA translation appears to be unique to Mena(Vidaki et al. 2017). Our detailed characterization of Mena protein-protein interactions in adult SNs, uncovered numerous interactions with translation-related molecules, as well as known interactions with components of the developing Mena-RNP(Vidaki et al. 2017). This points to a significant association of the protein with translation regulation that is either conserved in axons from development to adulthood, or reignited in response to injury, in order to facilitate regeneration.

In regeneration-potent neurons, injury triggers a series of dynamic events in two distinct temporal phases: an acute rapid response that aims to stabilize the injured axon tip and recreate a dynamic growth cone, while at the same time initiating retrograde transport of injury-signals to the soma, and a second slower phase that maintains retrograde signaling, prompts gene expression regulation and establishes anterograde transport of necessary materials to the axon, to support growth and regeneration(Batista and Hengst 2016; Smith et al. 2020; Mahar and Cavalli 2018). Both phases are highly dependent on axonal LT and appear to require Mena, according to our findings. Two hallmark proteins that are locally synthesized in axons of SNs upon injury, like Importin-β1(Perlson et al. 2004; Smith et al. 2020; Perlson et al. 2005) and mTOR(Terenzio et al. 2018), are significantly lower in Mena KO axons post-injury, contrary to naïve axons. Both proteins are essential for the acute response of axons to injury, and the initiation of the regenerative process. Importin-β1 is locally synthesized in axons of the PNS after axotomy, in order to bind phospho-ERK (extracellular-regulated kinase, ERK) and retrogradely transport it to the nucleus, for the activation of regeneration-associated genes (RAGs)(Perlson et al. 2004; Smith et al. 2020; Perlson et al. 2005). mTOR has also been shown to be locally translated after axotomy in the PNS, to modulate global translation and enhance regeneration(Terenzio et al. 2018). Notably, mTOR signaling contributes to both, the short-term activation of translation and to longer-term increases in the translational capacity of cells, thereby creating a positive feedback loop that sustains growth and regeneration(Wang and Proud 2006; Yang et al. 2022).

Interestingly, the mRNAs of *Kpnb1* (Importin-β1) and *mTOR*, do not appear to be specifically associated with Mena, according to a previous study that identified multiple cytosolic transcripts regulated by the Mena-RNP(Vidaki et al. 2017). Therefore, Mena could be modulating translation of *Kpnb1* and *mTOR* indirectly, via regulation of mTOR downstream signaling, which in turn regulates LT of numerous mRNAs, including that of *Kpnb1* and *mTOR*(Terenzio et al. 2018; Perry et al. 2016). However, one could not exclude the possibility that Mena may regulate additional mRNAs in adult axons, including *Kpnb1* and *mTOR*. This hypothesis could be tested in future studies. One of the best characterized substrates of mTOR is p70S6-Kinase (S6K1), which has been shown to be elevated as a result of LT in the PNS(Perry et al. 2016; Crino 2016). S6K1 subsequently phosphorylates ribosomal protein S6, to enhance protein synthesis(Batista and Hengst 2016; Terenzio et al. 2018; Yang et al. 2014), and eEF2-Kinase (eEF2K), leading to inhibition of eEF2 kinase activity, thus enhancing activity of eEF2 to promote translation elongation(Wang and Proud 2006; Wang et al. 2001). Our data demonstrate that all those crucial signaling effectors downstream of mTOR are significantly deregulated in Mena KO axons, disrupting the positive feedback loop of mTOR, and leading to significantly diminished eEF2 and S6 activation, as final effectors that mediate protein synthesis. This provides a plausible explanation as to the limited capacity of Mena-deficient nerves for LT and regeneration after injury.

One potential way that Mena functions to control mTOR activity and protein levels, is via regulating LT of the *Pik3r1* mRNA, thereby affecting the protein levels of PI3K-p85 regulatory subunit of PI3K. This, in turn, binds and stabilizes the PI3K-p110 catalytic subunit of the PI3K heterodimer, regulating its activity and downstream signaling in a cascade mode(Sánchez-Alegría et al. 2018; Taniguchi et al. 2010). Given the necessity of tightly regulated PI3K levels and activity to drive regeneration and functional recovery, even in the central nervous system (CNS)(Karova et al. 2025), the role of Mena could be pivotal for axon recovery. Appropriate levels and activation of PI3K are crucial to recruit downstream signaling proteins such as Akt(Guo et al. 2024; Sánchez-Alegría et al. 2018; Karova et al. 2025), with Akt1 being the most widely expressed subtype in neurons(Guo et al. 2024). According to our findings, Mena- deficiency not only affects protein levels of Akt1 after injury, but also perturbs its activity. A key effector downstream of Akt is mTOR, which forms two structurally and functionally distinct complexes. mTORC1 is directly regulated by Akt and controls cell growth, partly through its ability to phosphorylate S6K1 and other key regulators of mRNA translation(Guo et al. 2024; Dan et al. 2014). On the other hand, mTORC2 activity is PI3K-dependent and positively regulates Akt (S473), feeding back into mTORC1 regulation(Guo et al. 2024; Dan et al. 2014; Janku et al. 2018; Dai et al. 2014). We find that Mena deficiency disrupts the entire PI3K/Akt/mTOR signaling axis and any potential crosstalk.

In some cells ERK can also signal to activate mTOR(Hausott and Klimaschewski 2019; Dai et al. 2014; Ma et al. 2005). Interestingly, in the absence of Mena, we find phosphorylation of ERK significantly elevated, potentially as an attempt of the axons to restore mTOR activity; increased ERK phosphorylation could decrease the activity of eEF2K(via activation of p90 RSK1)(Wang et al. 2001; Guo et al. 2024), in order to balance the levels of eEF2 phosphorylation, and promote translation. Concomitantly, elevated ERK phosphorylation could increase injury-induced signaling to the nucleus, in order to sustain axon homeostasis and initiate regeneration after injury. In either case, this appears to be insufficient, as we demonstrate that Mena KO axons fail to increase mTOR levels and activity locally, as an acute response to injury *ex vivo*, which in turn leads to their inability to induce axonal protein synthesis and initiate a regenerative process. This essentially results in a diminished intrinsic capacity of Mena-deficient axons for regeneration after injury *in vivo*. Therefore, Mena-dependent LT of the PI3K regulatory subunit could function as an initial crucial step towards PI3K activation upon injury, coupling injury signals to the regenerative process.

Despite the pivotal role of LT in axonal response to injury and subsequent regeneration, its regulation in adult axons is not well understood. Our study provides new evidence on the modulation of acute axon responses to injury, and introduces a cytoskeleton-reorganizing protein as a key player that regulates the intrinsic ability of axons for LT and subsequent regeneration. Notably, the distinctive properties of Mena with respect to LT regulation and actin remodeling, render it an ideal candidate molecule to coordinate the two processes in axons, and potentially other cellular events downstream of different stimuli, including injury. Further future work is required to address the multifaceted roles of Mena in axon biology. However, our findings fit the notion that regeneration requires the engagement of developmental mechanisms in order to progress, and the translation-regulation role of Mena in adult axons may represent the reactivation of such mechanisms upon injury. Hence, understanding in depth the function of versatile molecules like Mena, could set the ground for therapeutic approaches in cases of degeneration or trauma in the central nervous system, where the inability of axons to regenerate poses a major challenge.

## MATERIALS & METHODS

### CONTACT FOR REAGENT AND RESOURCE SHARING

Further information and reagent requests may be directed to and will be fulfilled by Marina Vidaki

### Antibodies and Reagents

A full list of antibodies and reagents used in this study, is provided in the Supplemental Material

### Animals

All animal procedures were approved by the committee for Laboratory Animal Care at the University of Crete, and the Department of Welfare (Veterinarian administration; Ministry of Agriculture, Greece) (approval#69248). Experiments were conducted in accordance with the Greek National regulations on the use of vertebrate animals for experimental purposes, based on the ratification of the EU regulation for Laboratory Animal Use (86/609/CEE. Law N.1197/81ar.4 Law N.2015/92, Presidential Decree No 160/91ar. 12).

Adult mice (*Mus musculus*, strain C57BL/6J) (2-3 months old) of both sexes were used: wild-type (WT) C57BL/6J and Mena CRISPR/Cas9 (Mena KO) knock-out (C57BL/6J background, Taconic Biosciences). Animals were euthanized by cervical dislocation prior to tissue collection.

### *Ex vivo* Sciatic Nerve Injury Assay

Sciatic nerves of approximately 2 cm in length were harvested from adult wild-type and Mena KO mice. Nerves were subsequently sectioned into 4 equal segments, with each cutting point considered as an injury site. Segments were incubated in Dulbecco’s Modified Eagle Medium (DMEM; Gibco) supplemented with 10% fetal bovine serum (FBS; Gibco) and 1% penicillin-streptomycin (Gibco) at 37 °C in a 5% CO2 incubator for 2 and 4 hours respectively. Control segments were processed without any prior incubation (0 hours) to serve as a baseline condition.

### *In vivo* Sciatic Nerve Injury Assay

Mice were anesthetized with isoflurane (5% for anesthesia induction, 2.5% for anesthesia maintenance) and prepared for surgery by depilation of the hind limbs and lower back, followed by sterilization with povidone-iodine. An ophthalmic solution was applied to prevent corneal drying during surgery. A small incision was made to expose the sciatic nerve by blunt dissection of the biceps femoris and gluteus superficialis muscles. The nerve was crushed orthogonally for 20 seconds using 5-mm surgical forceps (91150-20, Inox Electronic), followed by carbon marking at the crush site. The crush was performed in a distance of approximately 20 mm from the nerve’s dorsal root ganglia (DRG).

### Dissociated DRG neuron culture

Neurons from 2MO mouse DRGs were cultured for 16 hrs in a humidified incubator (37°C, 5% CO2). Briefly, spinal cords were dissected and DRGs were removed in 10mM Hepes and 1x HBSS, washed in the same buffer and incubated with Collagenase IV (1mg/ml) for 30 min and Trypsin (0,1mg/ml) for another 15 min at 37°C. DRGs were then washed in DMEM 1x with 10% FBS to inactivate the enzymes, and triturated in the same medium. Following trituration neurons were pelleted at 600g for 5 min, resuspended in serum-free Neurobasal medium (Invitrogen), supplemented with B27 (Gibco), Pen/Strep (Gibco), and NGF (Nerve Growth Factor) (Gibco), and plated on coverslips that had been cleaned with 5 N hydrochloric acid and coated with 0.25 mg/ml poly-d-lysine (Sigma-Aldrich) and 100 μg/ml mouse Laminin 1 (Southern Biotech).

### Protein Extraction from Isolated Axoplasm

Sciatic nerve axoplasm was obtained by “light-squeezing” of the nerve segments in 200 μl of Axoplasm Isolation (AI) buffer (200 mM Tris, pH = 8.0; 2 mM EDTA) containing EDTA-free protease inhibitors (Millipore), via a glass Dounce Homogenizer. Total axoplasm proteins were obtained following centrifugation at 13.000 rpm for 15 minutes at 4°C and collection of supernatants. Supernatants were mixed with 4x Laemmli buffer (200mM Tris-HCl pH 6.8,8% Sodium Dodecyl sulfate, 40% Glycerol, 0.2% bromophenol blue, 0.4M DTT), followed by incubation at 95°C for 5 minutes, and used for SDS-PAGE and Western blot analysis.

### Sciatic Nerve Regeneration

Sciatic nerves were dissected 72 hours after surgery and post-fixed in 4% paraformaldehyde (PFA) at room temperature for 2 hours. The tissue was then transferred into 30% sucrose for three days. Subsequently, the tissue was embedded in a gelatin-sucrose solution, frozen, and stored at −80°C until sectioning. Sagittal tissue sections (12 µm thick) were obtained using a cryostat, and immunostained for GAP43 (1:500, Abcam) as a marker of nerve regeneration. The crush site was identified by deformation of the nerve and disruption of axons, which coincided with the highest GAP43 intensity. Four sections per animal and at least four biological replicates were analyzed. Imaging was performed using a Leica Sp8 Confocal Microscope at 10x magnification.

### Puromycin Incorporation

Sciatic nerve segments from wild-type and Mena-deficient mice were incubated in Dulbecco’s Modified Eagle Medium (DMEM) supplemented with 10% fetal bovine serum (FBS) and 1% penicillin/streptomycin at 37°C with 5% CO₂ for 4 hours. For experimental conditions, 50 µg/mL puromycin was added to the medium. Control conditions included sciatic nerve segments without puromycin and non-incubated sciatic nerves (0-hour control). Following incubation, sciatic nerve segments were post-fixed in 4% paraformaldehyde (PFA) at room temperature for 2 hours and transferred to 30% sucrose for three days. The tissue was embedded in a gelatin-sucrose solution, frozen, and stored at −80°C until sectioning. Tissue sections (12 µm thick) were prepared in both sagittal and coronal planes. Sections were immunostained for Puromycin (1:500, Millipore), and Tubulin (1:200, Chemicon). Imaging was performed using a Leica Sp8 Confocal Microscope at 42× magnification. At least three biological replicates were analyzed, with a minimum of four sections per animal examined.

A detailed description of additional methods used in this study is provided in the Supplemental Material.

## AKNOWLEDGEMENTS

The authors would like to thank M. Denaxa, F. Gertler, D. Karagogeos, Z. Kontarakis, M. Savvaki, A. Tatarakis, E. Trompouki and N. Mizuno for helpful discussion on the manuscript.

This work was supported by the Hellenic Foundation for Research and Innovation (HFRI, grants 1442 and 2343, MV), and funds from the Fondation Sante (MV)

## AUTHOR CONTRIBUTIONS

SP designed and performed experiments, analyzed data and assisted with writing the manuscript. NT performed bioinformatic analysis, developed the axoplasm preparation protocol and assisted with writing the manuscript. CK and ET performed *in vitro* DRG neuron cultures. EK performed WB assays. MS performed Mass Spectrometry and initial analysis of proteomic data. MV designed and supervised experiments, analyzed data and wrote the manuscript.

## Abbreviations

LT: Local translation
RNP: Ribonucleoprotein complex
RBP: RNA-binding Proteins
PNS: Peripheral Nervous System
SN: Sciatic Nerve
SNC: Sciatic Nerve Crush

## REFERENCES

Akram R, Anwar H, Javed MS, Rasul A, Imran A, Malik SA, Raza C, Khan IU, Sajid F, Iman T, et al. 2022. Axonal Regeneration: Underlying Molecular Mechanisms and Potential Therapeutic Targets. Biomedicines 10: 3186. https://www.mdpi.com/2227-9059/10/12/3186 (Accessed October 16, 2024).

Al-Ali H, Ding Y, Slepak T, Wu W, Sun Y, Martinez Y, Xu X-M, Lemmon VP, Bixby JL. 2017. The mTOR Substrate S6 Kinase 1 (S6K1) Is a Negative Regulator of Axon Regeneration and a Potential Drug Target for Central Nervous System Injury. J Neurosci 37: 7079–7095. https://www.jneurosci.org/lookup/doi/10.1523/JNEUROSCI.0931-17.2017 (Accessed October 18, 2024).

Altas B, Romanowski AJ, Bunce GW, Poulopoulos A. 2022. Neuronal mTOR Outposts: Implications for Translation, Signaling, and Plasticity. Front Cell Neurosci 16: 853634. https://www.ncbi.nlm.nih.gov/pmc/articles/PMC9021820/ (Accessed July 26, 2024).

Baleriola J, Walker CA, Jean YY, Crary JF, Troy CM, Nagy PL, Hengst U. 2014. Axonally synthesized ATF4 transmits a neurodegenerative signal across brain regions. Cell 158: 1159–1172.

Barzik M, Kotova TI, Higgs HN, Hazelwood L, Hanein D, Gertler FB, Schafer DA. 2005. Ena/VASP proteins enhance actin polymerization in the presence of barbed end capping proteins. J Biol Chem 280: 28653–28662.

Bashaw GJ, Kidd T, Murray D, Pawson T, Goodman CS. 2000. Repulsive axon guidance: Abelson and Enabled play opposing roles downstream of the roundabout receptor. Cell 101: 703–715.

Batista AFR, Hengst U. 2016. Intra-axonal protein synthesis in development and beyond. Int J Dev Neurosci 55: 140–149.

Bear JE, Svitkina TM, Krause M, Schafer DA, Loureiro JJ, Strasser GA, Maly IV, Chaga OY, Cooper JA, Borisy GG, et al. 2002. Antagonism between Ena/VASP proteins and actin filament capping regulates fibroblast motility. Cell 109: 509–521.

Bradke F, Fawcett JW, Spira ME. 2012. Assembly of a new growth cone after axotomy: the precursor to axon regeneration. Nat Rev Neurosci 13: 183–193. http://www.nature.com/articles/nrn3176 (Accessed January 17, 2022).

Christie KJ, Webber CA, Martinez JA, Singh B, Zochodne DW. 2010. PTEN inhibition to facilitate intrinsic regenerative outgrowth of adult peripheral axons. J Neurosci 30: 9306–9315.

Crino PB. 2016. The mTOR signalling cascade: paving new roads to cure neurological disease. Nat Rev Neurol 12: 379–392. https://www.nature.com/articles/nrneurol.2016.81 (Accessed November 13, 2024).

Dai J, Bercury KK, Macklin WB. 2014. Interaction of mTOR and Erk1/2 signaling to regulate oligodendrocyte differentiation. Glia 62: 2096–2109. https://onlinelibrary.wiley.com/doi/abs/10.1002/glia.22729 (Accessed January 8, 2025).

Dan HC, Ebbs A, Pasparakis M, Dyke TV, Basseres DS, Baldwin AS. 2014. Akt-dependent Activation of mTORC1 Complex Involves Phosphorylation of mTOR (Mammalian Target of Rapamycin) by IκB Kinase α (IKKα). The Journal of Biological Chemistry 289: 25227. https://pmc.ncbi.nlm.nih.gov/articles/PMC4155685/ (Accessed January 12, 2025).

Dent EW, Gupton SL, Gertler FB. 2011. The Growth Cone Cytoskeleton in Axon Outgrowth and Guidance. Cold Spring Harbor Perspectives in Biology 3: a001800–a001800. http://cshperspectives.cshlp.org/lookup/doi/10.1101/cshperspect.a001800 (Accessed January 17, 2022).

Donnelly CJ, Willis DE, Xu M, Tep C, Jiang C, Yoo S, Schanen NC, Kirn-Safran CB, van Minnen J, English A, et al. 2011. Limited availability of ZBP1 restricts axonal mRNA localization and nerve regeneration capacity. EMBO J 30: 4665–4677.

Drees F, Gertler FB. 2008. Ena/VASP: proteins at the tip of the nervous system. Curr Opin Neurobiol 18: 53–59.

Gebauer F, Hentze MW. 2004. Molecular mechanisms of translational control. Nat Rev Mol Cell Biol 5: 827–835.

Gertler FB, Niebuhr K, Reinhard M, Wehland J, Soriano P. 1996. Mena, a relative of VASP and Drosophila Enabled, is implicated in the control of microfilament dynamics. Cell 87: 227–239.

Guo N, Wang X, Xu M, Bai J, Yu H, Le Zhang. 2024. PI3K/AKT signaling pathway: Molecular mechanisms and therapeutic potential in depression. Pharmacological Research 206: 107300. https://www.sciencedirect.com/science/article/pii/S1043661824002457 (Accessed January 12, 2025).

Gupton SL, Riquelme D, Hughes-Alford SK, Tadros J, Rudina SS, Hynes RO, Lauffenburger D, Gertler FB. 2012. Mena binds α5 integrin directly and modulates α5β1 function. J Cell Biol 198: 657–676.

Hausott B, Klimaschewski L. 2019. Promotion of Peripheral Nerve Regeneration by Stimulation of the Extracellular Signal-Regulated Kinase (ERK) Pathway. The Anatomical Record 302: 1261–1267. https://onlinelibrary.wiley.com/doi/abs/10.1002/ar.24126 (Accessed January 8, 2025).

Janku F, Yap TA, Meric-Bernstam F. 2018. Targeting the PI3K pathway in cancer: are we making headway? Nat Rev Clin Oncol 15: 273–291. https://www.nature.com/articles/nrclinonc.2018.28 (Accessed January 12, 2025).

Jung H, Yoon BC, Holt CE. 2012. Axonal mRNA localization and local protein synthesis in nervous system assembly, maintenance and repair. Nat Rev Neurosci 13: 308–324. https://www.nature.com/articles/nrn3210 (Accessed June 17, 2024).

Karova K, Polcanova Z, Knight L, Suchankova S, Nieuwenhuis B, Holota R, Herynek V, Machova Urdzikova L, Turecek R, Kwok JC, et al. 2025. Hyperactive delta isoform of PI3 kinase enables long-distance regeneration of adult rat corticospinal tract. Mol Ther 33: 752–770. https://pmc.ncbi.nlm.nih.gov/articles/PMC11852985/ (Accessed September 29, 2025).

Krause M, Dent EW, Bear JE, Loureiro JJ, Gertler FB. 2003. Ena/VASP proteins: regulators of the actin cytoskeleton and cell migration. Annu Rev Cell Dev Biol 19: 541–564.

Kwiatkowski AV, Rubinson DA, Dent EW, Edward van Veen J, Leslie JD, Zhang J, Mebane LM, Philippar U, Pinheiro EM, Burds AA, et al. 2007. Ena/VASP Is Required for neuritogenesis in the developing cortex. Neuron 56: 441–455.

Lanier LM, Gates MA, Witke W, Menzies AS, Wehman AM, Macklis JD, Kwiatkowski D, Soriano P, Gertler FB. 1999. Mena Is Required for Neurulation and Commissure Formation. Neuron 22: 313–325. https://linkinghub.elsevier.com/retrieve/pii/S0896627300810922 (Accessed January 17, 2022).

Lebrand C, Dent EW, Strasser GA, Lanier LM, Krause M, Svitkina TM, Borisy GG, Gertler FB. 2004. Critical role of Ena/VASP proteins for filopodia formation in neurons and in function downstream of netrin-1. Neuron 42: 37–49.

Ma L, Chen Z, Erdjument-Bromage H, Tempst P, Pandolfi PP. 2005. Phosphorylation and functional inactivation of TSC2 by Erk implications for tuberous sclerosis and cancer pathogenesis. Cell 121: 179–193.

Mahar M, Cavalli V. 2018. Intrinsic mechanisms of neuronal axon regeneration. Nature reviews Neuroscience 19: 323. https://pmc.ncbi.nlm.nih.gov/articles/PMC5987780/ (Accessed January 9, 2025).

McConnell RE, Edward van Veen J, Vidaki M, Kwiatkowski AV, Meyer AS, Gertler FB. 2016. A requirement for filopodia extension toward Slit during Robo-mediated axon repulsion. J Cell Biol 213: 261–274.

Menzies AS. 2004. Mena and Vasodilator-Stimulated Phosphoprotein Are Required for Multiple Actin-Dependent Processes That Shape the Vertebrate Nervous System. Journal of Neuroscience 24: 8029–8038. https://www.jneurosci.org/lookup/doi/10.1523/JNEUROSCI.1057-04.2004 (Accessed January 17, 2022).

Nagano S, Araki T. 2021. Axonal Transport and Local Translation of mRNA in Neurodegenerative Diseases. Frontiers in Molecular Neuroscience 14: 697973. https://pmc.ncbi.nlm.nih.gov/articles/PMC8236635/ (Accessed October 21, 2024).

Perlson E, Hanz S, Ben-Yaakov K, Segal-Ruder Y, Seger R, Fainzilber M. 2005. Vimentin-Dependent Spatial Translocation of an Activated MAP Kinase in Injured Nerve. Neuron 45: 715–726. https://linkinghub.elsevier.com/retrieve/pii/S0896627305000553 (Accessed July 3, 2024).

Perlson E, Medzihradszky KF, Darula Z, Munno DW, Syed NI, Burlingame AL, Fainzilber M. 2004. Differential proteomics reveals multiple components in retrogradely transported axoplasm after nerve injury. Mol Cell Proteomics 3: 510–520.

Perry RB-T, Rishal I, Doron-Mandel E, Kalinski AL, Medzihradszky KF, Terenzio M, Alber S, Koley S, Lin A, Rozenbaum M, et al. 2016. Nucleolin-Mediated RNA Localization Regulates Neuron Growth and Cycling Cell Size. Cell Rep 16: 1664–1676. https://www.ncbi.nlm.nih.gov/pmc/articles/PMC4978702/ (Accessed August 8, 2024).

Rishal I, Rozenbaum M, Fainzilber M. 2010. Axoplasm isolation from rat sciatic nerve. J Vis Exp 2087.

Sánchez-Alegría K, Flores-León M, Avila-Muñoz E, Rodríguez-Corona N, Arias C. 2018. PI3K Signaling in Neurons: A Central Node for the Control of Multiple Functions. International Journal of Molecular Sciences 19: 3725. https://pmc.ncbi.nlm.nih.gov/articles/PMC6321294/ (Accessed January 8, 2025).

Saxton RA, Sabatini DM. 2017. mTOR Signaling in Growth, Metabolism, and Disease. Cell 168: 960–976.

Serger E, Luengo-Gutierrez L, Chadwick JS, Kong G, Zhou L, Crawford G, Danzi MC, Myridakis A, Brandis A, Bello AT, et al. 2022. The gut metabolite indole-3 propionate promotes nerve regeneration and repair. Nature 607: 585–592. https://www.nature.com/articles/s41586-022-04884-x (Accessed November 12, 2024).

Shannon P, Markiel A, Ozier O, Baliga NS, Wang JT, Ramage D, Amin N, Schwikowski B, Ideker T. 2003. Cytoscape: A Software Environment for Integrated Models of Biomolecular Interaction Networks. Genome Res 13: 2498–2504. https://genome.cshlp.org/content/13/11/2498 (Accessed July 24, 2024).

Smith TP, Sahoo PK, Kar AN, Twiss JL. 2020. Intra-axonal mechanisms driving axon regeneration. Brain Research 1740: 146864. https://linkinghub.elsevier.com/retrieve/pii/S0006899320302201 (Accessed January 17, 2022).

Taniguchi CM, Winnay J, Kondo T, Bronson RT, Guimaraes AR, Aleman JO, Luo J, Stephanopoulos G, Weissleder R, Cantley LC, et al. 2010. The PI3K regulatory subunit p85α can exert tumor suppressor properties through negative regulation of growth factor signalling. Cancer research 70: 5305. https://pmc.ncbi.nlm.nih.gov/articles/PMC3204358/ (Accessed January 8, 2025).

Terenzio M, Koley S, Samra N, Rishal I, Zhao Q, Sahoo PK, Urisman A, Marvaldi L, Oses-Prieto JA, Forester C, et al. 2018. Locally translated mTOR controls axonal local translation in nerve injury. Science 359: 1416–1421.

Triantopoulou N, Vidaki M. 2022. Local mRNA translation and cytoskeletal reorganization: Mechanisms that tune neuronal responses. Front Mol Neurosci 15: 949096.

Van Dongen S. 2008. Graph Clustering Via a Discrete Uncoupling Process. SIAM J Matrix Anal Appl 30: 121–141. https://epubs.siam.org/doi/10.1137/040608635 (Accessed January 27, 2025).

Verma P, Chierzi S, Codd AM, Campbell DS, Meyer RL, Holt CE, Fawcett JW. 2005. Axonal Protein Synthesis and Degradation Are Necessary for Efficient Growth Cone Regeneration. J Neurosci 25: 331–342. https://www.ncbi.nlm.nih.gov/pmc/articles/PMC3687202/ (Accessed June 14, 2024).

Vidaki M, Drees F, Saxena T, Lanslots E, Taliaferro MJ, Tatarakis A, Burge CB, Wang ET, Gertler FB. 2017. A Requirement for Mena, an Actin Regulator, in Local mRNA Translation in Developing Neurons. Neuron 95: 608–622.e5.

Wang X, Li W, Williams M, Terada N, Alessi DR, Proud CG. 2001. Regulation of elongation factor 2 kinase by p90RSK1 and p70 S6 kinase. The EMBO Journal 20: 4370–4379. https://www.embopress.org/doi/full/10.1093/emboj/20.16.4370 (Accessed January 10, 2025).

Wang X, Proud CG. 2006. The mTOR Pathway in the Control of Protein Synthesis. Physiology 21: 362–369. https://www.physiology.org/doi/10.1152/physiol.00024.2006 (Accessed January 10, 2025).

Yang L, Miao L, Liang F, Huang H, Teng X, Li S, Nuriddinov J, Selzer ME, Hu Y. 2014. The mTORC1 effectors S6K1 and 4E-BP play different roles in CNS axon regeneration. Nat Commun 5: 5416. https://www.nature.com/articles/ncomms6416 (Accessed August 5, 2024).

Yang M, Lu Y, Piao W, Jin H. 2022. The Translational Regulation in mTOR Pathway. Biomolecules 12: 802.

Yu G, Wang L-G, Han Y, He Q-Y. 2012. clusterProfiler: an R package for comparing biological themes among gene clusters. OMICS 16: 284–287.

